# The HIV-1 Integrase C-Terminal domain induces TAR RNA structural changes promoting Tat binding

**DOI:** 10.1101/2021.10.21.465253

**Authors:** Cecilia Rocchi, Camille Louvat, Adriana Miele, Julien Batisse, Christophe Guillon, Lionel Ballut, Daniela Lener, Matteo Negroni, Marc Ruff, Patrice Gouet, Francesca Fiorini

**Author notes:** equal contribution (Cecilia Rocchi, Camille Louvat).

## Abstract

Recent evidence indicated that HIV-1 Integrase (IN) binds genomic viral RNA (gRNA) playing a critical role in viral particle morphogenesis and gRNA stability in host cells. Combining biophysical and biochemical approaches we show that the C-terminal flexible 18-residues tail of IN acts as a sensor of the peculiar apical structure of trans-activation response element RNA (TAR), directly interacting with its hexaloop. We highlighted how the whole IN C-terminal domain, once bound to TAR, can change its structure assisting the binding of Tat, the HIV trans-activator protein, which finally displaces IN from TAR. Our results are consistent with the emerging role of IN in early stage of proviral transcription and suggest new steps of HIV-1 life cycle that can be considered as therapeutic targets.

## INTRODUCTION

Protein-nucleic acid interactions can occur through different types of protein binding domains and are responsible for a variety of essential molecular and cellular mechanisms and their regulation. This binding diversity has been well described by recent RNA interactome screenings revealing that the term RNA-Binding Domain (RBD) is no longer synonymous with a well-structured domain, but also with intrinsically disordered region (IDR) with non-canonical RNA-binding properties ^1 2 3^. Moreover, an increasing level of complexity has been documented for certain transcription factors that are able to bind both DNA and RNA through separated structured or unstructured regions, resulting in a complex pattern of specific and non-specific interactions ^4 5^. This protein moonlighting is particularly true for RNA viruses which possess a relatively short genome encoding only for a small amount of proteins that must ensure many multiple functions during viral replication ^6 7^.

A good example of moonlighting is represented by Human Immunodeficiency Virus type 1 (HIV-1) integrase (IN) which can bind both DNA and RNA ^8,9^. As for all retroviruses, IN catalyzes the integration of viral cDNA (vDNA), produced by retro-transcription of genomic RNA (gRNA), into the host genome. Multimeric IN binds the vDNA ends forming the intasome complex able to catalyze the processing of 3’-end dinucleotides. After activation, the 3’ extremities are then used by the intasome to attack the host DNA in order to integrate the provirus (reviewed in ^9^). A step essential for HIV-1 productive infection ^10^. Retroviral integrases are modular proteins that contain three structured domains: the N-terminal domain (NTD), the catalytic core domain (CCD) and the C-terminal domain (CTD), connected by unstructured regions. All three domains show protein-protein and protein-DNA interaction properties and are essential for enzymatic activity (reviewed in ^9^and ^11^). The CCD harbors the essential catalytic triad D, D, E that coordinates two Mg^2+^ cofactors and folds similarly to nucleotidyl-transferases and nucleases ^12^. The NTD is involved in enzyme multimerization and catalytic activity and shows a HHCC motif coordinating a Zn^2+^ ion ^13 14 15^. The CTD is also involved in DNA interaction, multmerization, and possesses a SH3-like fold followed by a flexible 18-residues tail (CT) ^16 17 18 19^. The SH3-like domain of IN is the minimal DNA binding site. In HIV-1 life cycle, this hub domain mediates the interaction with RT, with cellular nuclear import complex TRN-SR2; and with histone 4 tail, likely anchoring the intasome to the chromatin and therefore promoting an efficient integration ^20 21 22 23^. The mutational study of flexible CT revealed a moderate implication in IN enzymatic activity ^21 24^, but significant effect on the incorporation of IN in virions and on HIV-1 infectivity ^21^. However the exact function of CT region remains largely unknown.

As mentioned before, recent works revealed that in HIV-1, IN is also an RBP with an essential role in virion morphogenesis, related to its ability to bind gRNA ^8 25 26^. In fact, IN interacts with specific sites within gRNA ensuring the correct localization of viral RNP inside the capsid (reviewed in ^27^). Aberrant virions are obtained when IN-gRNA interaction is abolished, highlighting the importance of the proper formation of IN-containing RNPs for HIV-1 infection (^8,25,28^). In addition, the IN-gRNA interaction is also dictating the fate of the gRNA within the host cell in the early steps of infection. Indeed, when virions are defective for IN-gRNA interactions, viral infection is blocked at early stages of reverse transcription, due to a rapid degradation of gRNA in host cells ^25,28^. If the RBD within IN has not yet been structurally identified, most of the Lysine residues interacting with RNA are located within the CTD and overlap with those subjected to post-translational modification ^8 29 30 31^. Importantly, Lysine 273, belonging to the flexible CT, seems to be the only Lysine dedicated to RNA interaction to be essential for viral infectivity ^8^.

Recently, a new role has been proposed for HIV-1 IN during proviral transcription at early times after integration. In fact, after strand transfer, the IN remains bound to DNA and directly plays a role in proviral transcription, depending on its post-translational modifications of specific residues within the CTD ^32^.

HIV-1 provirus is transcribed by the cellular RNA polymerase II (Pol II) which pauses shortly after initiation of transcription, due to the presence of negative elongation factors as well as nucleosomes downstream the transcription start site ^33 34^. HIV-1 removes this block by encoding a transcriptional trans-activator Tat protein, which binds the nascent transcript on a structured RNA sequence named TAR (trans-activation response element RNA) using a non-canonical RBD. This allows the recruitment of the human super elongation complex (SEC) ^35 36 37 38^. In particular, Tat binds to p-TEFb, a complex composed of CDK9 kinase and its regulatory partner the cyclin T1 (CycT1), and consequently drives SEC to TAR RNA. This complex triggers a cascade of phosphorylation of several transcription factors, which activate Pol II and recruit positive chromatin remodelers. Moreover, processivity of Pol II is also enhanced by pTEFb-mediated phosphorylation of its C-terminal domain ^39^ (reviewed in ^40^). Unfortunately, molecular details about the interplay between IN and this cellular transcription initiation machinery are largely unknown so far.

Crosslinking-immuno-precipitation sequencing (CLIP-seq) experiments have identified the TAR RNA sequence as a major binding site of HIV-1 IN on the gRNA ^8^. This observation, together with the recent finding of HIV-1 IN involvement in proviral transcription ^32^, prompted us to study the interaction of IN with TAR RNA and its interplay with the Tat protein. Our results revealed that despite the apparent lack of structural specificity of IN *in vitro*, the CT flexible tail discriminates for the proper TAR apical stem-loop. We described the consequences of the IN binding on the structure of TAR and on the subsequent Tat/TAR interaction, proposing a working model which foresees a possible involvement of IN in proviral transcription elongation before the arrival of Tat.

## RESULTS

### IN binds TAR RNA with no apparent structural specificity

We first addressed whether full-length HIV-1 IN was able to specifically bind TAR RNA. One hindering aspect of this study is the well-known low solubility of HIV-1 IN as well as its flexibility between N-terminal (NTD), C-terminal (CTD) and catalytic core (CCD) domains that for long time frustrated structural studies (**Fig. 1a** ^41^). The poor solubility *in vitro* is usually overcome by mutations of hydrophobic residues, however resulting in a replication-defective virus with mislocalized viral RNP phenotype analogous to that observed in IN mutants defective for the IN-gRNA interaction ^19 8,25,28^. We have chosen to express recombinant wild-type N-Terminal Flag-tagged IN-FL in eukaryotic expression system as previously described (^42^ **Supplementary Fig. 1a**, left panel) and called it IN-FLm (1-288 amino acids [aa]). Mass spectrometry analysis of this protein revealed post-translational modification at several residues: Serine 24 was phosphorylated and Lysine 46, 173, 211 and 273 residues were acetylated. The protein produced in mammalian cells was shown to have an increased solubility as compared to that produced in *E. coli* and also an enhanced enzymatic activity *in vitro* ^42^. However, after purification, we kept the protein in solution after buffer optimization. In this study we used an electrophoretic mobility shift assay (EMSA) in order to assess qualitatively the binding of IN to a synthetic TAR RNA, whose secondary structure is depicted in **Fig. 1b**, to a weakly structured RNA_(30)_-mer (**Supplementary Fig. 1b**, left panel) and to an unstructured AG_(50)_-mer RNA (**Fig. 1c**). We have incubated radiolabeled RNA with increasing concentrations of purified IN-FLm. A band shift was observed with TAR RNA upon gel electrophoresis under nearly physiological salt concentration, reflecting the formation of an IN:TAR complex (**Fig. 1c**). IN-FLm also bound weakly-structured RNA_(30)_-mer (**Supplementary Fig. 1b**, right panel), while it had a markedly reduced affinity for unstructured AG_(50)_-mer RNA (**Fig. 1c**). IN-FL produced in prokaryotic expression system (**Supplementary Fig. 1a**, right panel) is also able to bind TAR (**Supplementary Fig. 1c**).

**Figure 1:**
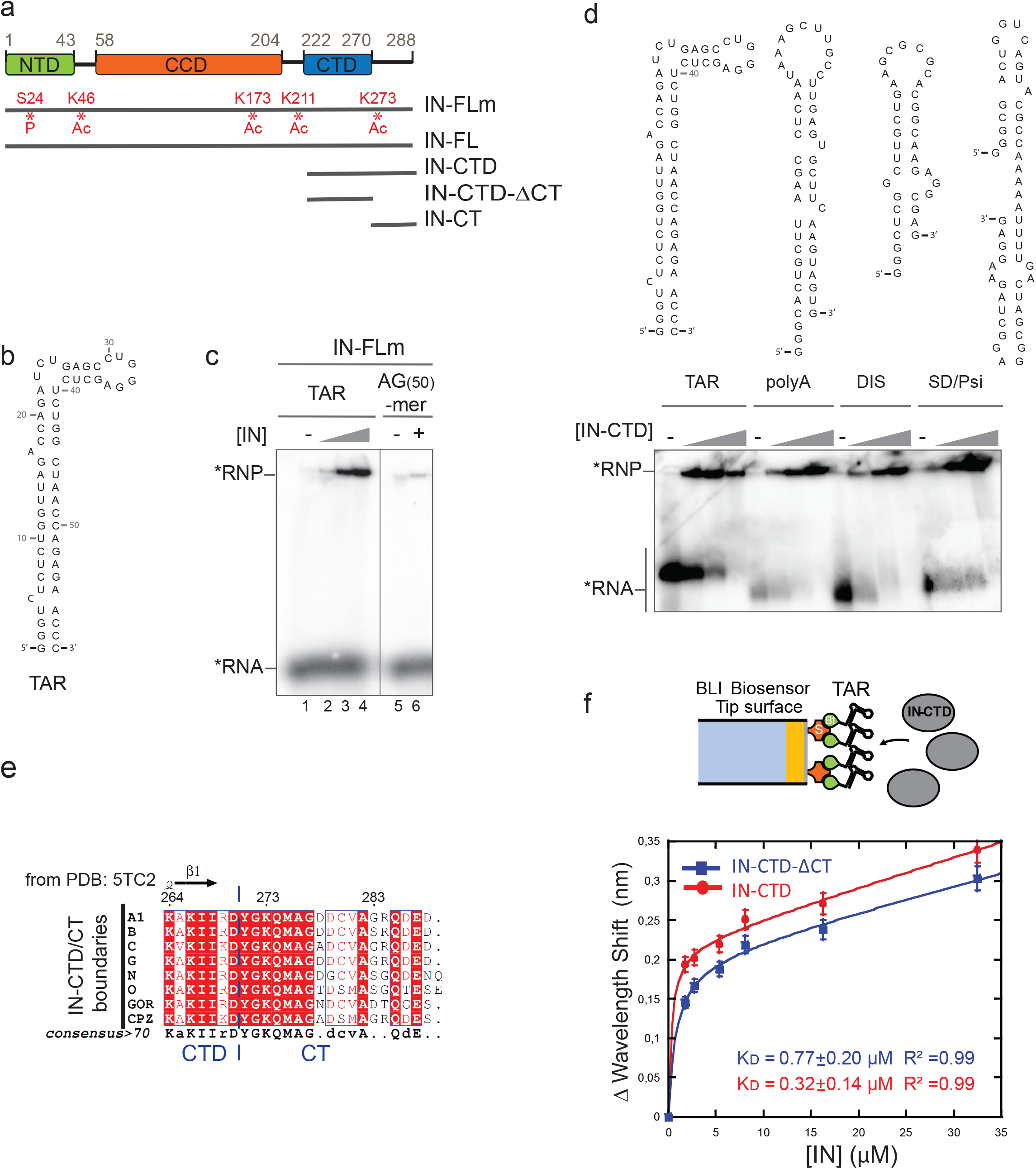
IN binds to structured RNA. **a** Schematic diagram showing the domain organization of IN: the N-terminal domain (NTD), the catalytic core domain (CCD) and the C-terminal domain (CTD) are indicated respectively in green, orange and blue rectangles. Recombinant protein versions used in our study are represented by grey lines: IN full-length from mammalian expression system (IN-FLm); IN full-length from *E. coli* (IN-FL); IN-CTD (residues 222 to 288); IN-CTD-ΔCT (222 to 270); IN-CT (270 to 288). Sites of phosphorylation (P) and acetylation (Ac) are shown in red. **b** Structural model of TAR RNA obtained with RNA Fold WebServer ^63 64^ predicting a minimum free energy of −29.60 kcal/mol. **c** Representative native 6% polyacrylamide gel illustrating the Electrophoretic mobility shift assay (EMSA) showing the interaction of IN-FLm with TAR RNA or an unstructured RNA 50-mer (AG_(50)_-mer) labelled with ^32^P (black star). The RNA substrates (50 nM) were incubated with various concentrations of IN-FLm or without protein (TAR RNA: 0; 100, 200, 400 nM of IN-FLm; AG_(50)_-mer RNA: 0 or 400 nM of IN-FLm). **d** Models of secondary structures of four RNA elements belonging to 5’ untranslated region of the HIV-1 RNA genome: TAR, Poly-A, DIS and SD/Psi (upper panel). EMSA assay showing the binding of IN-CTD to these RNA elements (lower panel). Increasing amounts of IN-CTD (0; 100; 200; 400 nM) were incubated with 5’-end radiolabeled RNA (50nM). **e** Sequence alignment of the C-terminal extremity of IN-CTD from HIV-1 subtypes and simian viruses. All amino acid sequences were obtained from the HIV database compendium (http://www.hiv.lanl.gov/) and aligned using Clustal Omega (https://www.ebi.ac.uk/Tools/msa/clustalo/) in order to have a consensus sequence for each viral subtype. Consensus sequences from subtypes A1, B, C, G, N, O and from GOR and CPZ, where aligned and analyzed by ESPript 3.0 Web server ^65^. Secondary structure elements from IN-CTD-ΔCT structure (PDB code: 5TC2) were presented on top of the alignment (helices with squiggles and strands with arrows). Red shading indicates sequence identity and boxes indicate sequence similarity, according to physical-chemical properties. **f** top: Schematic representation of the Bio-Layer interferometry (BLI) experiment showing the binding of IN (grey ellipses) to 5’-biotinylated TAR RNA immobilized on streptavidin-coated biosensor; bottom: graph showing the wavelength shifts recorded at 200 s after the start of the protein/RNA binding were plotted against the corresponding IN-CTD-ΔCT (blue line, squares) and IN-CTD (red line, circles) concentrations, in order to calculate the respective equilibrium dissociation constant (K_D_) values. Data points were fitted to the equation: Y = B_max_ * X/(Kd+X)+NS*X where B_max_ is the maximum wavelength shift and NS the slope of the non-linear component as described in ^66^. The coefficients of determination (R^2^) and equilibrium dissociation constant (K_D_) values obtained are indicated in the graph for each IN fragment. Binding assays were performed in duplicate. Error bars indicate the Standard Error of the Mean.

As RNA-interacting lysine residues are contained within the C-terminal domain of IN ^8^, we focused on this domain. We expressed and purified the whole C-Terminal Domain (IN-CTD; aa 222-288). We first determined whether IN-CTD was able to interact with structurally distinct viral genomic RNA elements of similar nucleotide length derived from HIV-1 5′ UTR. In addition to TAR, we synthesized three other RNA hairpins: the polyadenylation (polyA) signal, the dimerization initiation sequence (DIS) and the major splice-donor (SD) together with Ψ packaging element (Psi) (**Fig. 1d**, top panel). IN-CTD bound all those RNAs with no measurable difference (**Fig. 1d**, bottom panel), suggesting that IN can bind structured RNA through its C-terminal domain.

Subsequently, we wanted to address whether the C-terminal flexible tail (CT) spanning the last 18 residues of CTD affected the RNA binding properties of IN. We expressed the CTD without its terminal tail (IN-CTD-ΔCT; aa 220-270, **Fig. 1a**) and used a chemically synthesized IN-CT peptide (aa 270-288; **Fig. 1a**). The boundaries between CTD and CT were defined according to sequence alignment and previous structural studies ^16 43,44 45^. The IN-CT is not conserved among lentiviruses (**Supplementary Fig. 2a**), however, multiple alignment of IN-CT from human HIV-1 subtypes and simian viruses (**Fig. 1e**) showed a 58 % of sequence identity and 87 % of similarity (equivalent residues considering physical-chemical properties).

EMSA assays did not shown apparent differences between the affinities of IN-CTD and IN-CTD-ΔCT for TAR under physiological salt conditions (**Supplementary Fig. 2b**). To measure the rate constants of the IN-CTD and IN-CTD-ΔCT interactions with TAR, we used Bio-layer interferometry (BLI). A 3’ biotinylated TAR RNA, was immobilized on a streptavidin-coated biosensor to serve as a bait molecule (**Fig. 1f**, top panel). Subsequently, the interactions of IN subdomains with the TAR RNA were monitored in real time and the resulting sensorgrams are shown in **Supplementary Fig. 2c**. Fitted data resulted in K_D_ values of 0.77 and 0.32 μM for IN-CTD-ΔCT and IN-CTD, respectively (**Fig. 1f**, bottom panel). Thus, the presence of the 18 aa C-terminal tail only slightly affects the RNA binding affinity of IN-CTD.

### The C-terminal Tail senses the TAR RNA shape

It has been shown that the presence of the bulge and the loop in TAR RNA, rather than its sequence, is critical for IN binding, as the deletion of one or both markedly decreased IN binding affinity ^8^. We wondered whether the shape of the peculiar apical stem-loop of TAR (**Fig. 2a**) could also be important for IN binding. Therefore, we mutated the 4-nucleotides (nt) stem between the bulge and the loop to change its length, as shortening or lengthening the 4 nt-stem is likely to tighten or loosen the TAR major groove by bringing the bulge and the hexa-loop closer or further, respectively. We produced TAR with 1 bp longer stem (TAR-LS) and two with progressively shorter stem TARs (TAR-SS and TAR-VSS) (**Fig. 2a**). The EMSA assay showed that IN-FLm was able to bind TAR-LS similarly to the wild-type TAR (**Supplementary Fig. 3a**, lines A and B). On the contrary, shortening of the stem reduced IN-FLm affinity for TAR (**Supplementary Fig. 3a**, lines C to E). Furthermore, we assessed the binding behavior of IN-CTD and INCTD-ΔCT to TAR mutants by EMSA (**Fig. 2b** and **2c**). As observed for the full-length protein, the affinity of IN-CTD for TAR mutants decreased as the length of the stem was reduced (**Fig. 2b**, lines A to D and **Fig. 2c**). Surprisingly, the binding ability of IN-CTD-ΔCT was not affected by the shortening of the TAR 4nt-stem (**Fig. 2b**, lines A’ to D’, and **Fig. 2c**), suggesting that IN C-terminal tail senses the shape of the TAR RNA stem *in vitro*.

**Figure 2.**
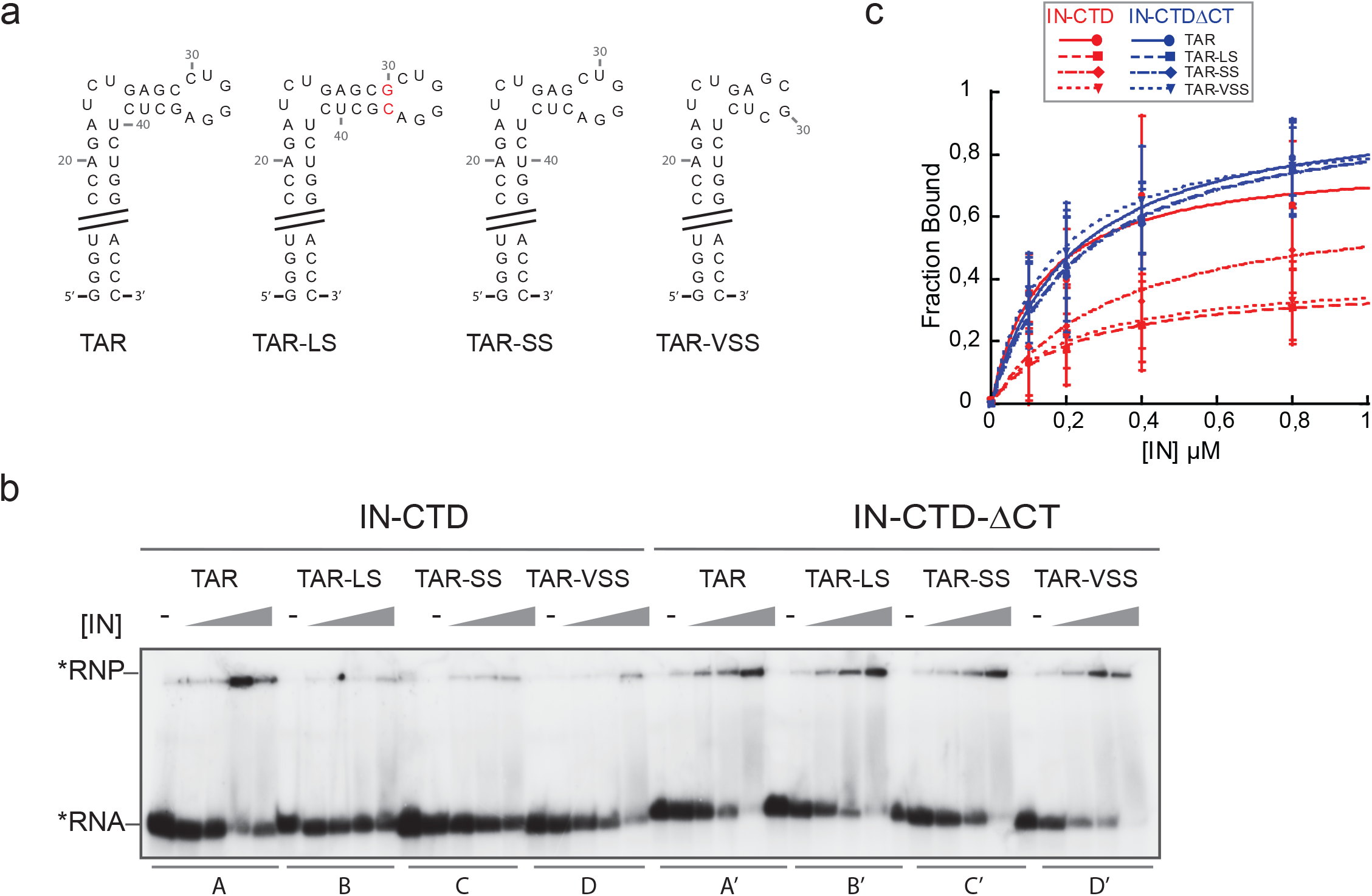
IN C-terminal tail sensing for TAR RNA apical stem-loop shape. **a** Structural model of TAR RNA and mutated versions used in our study: TAR long stem (TAR-LS), TAR short stem (TAR-SS) and TAR very short stem (TAR-VSS). In order to verify that each RNA mutant assumed the expected secondary structure each model was analyzed with Fold WebServer ^63,64^ **b** EMSA assay illustrating the interaction of IN-CTD and IN-CTD-ΔCT with TAR wild type and mutants. The RNA substrates (50 nM) are labelled with ^32^P (black star) and incubated with increasing concentrations of proteins (0; 100, 200, 400 and 800 nM) under the conditions described in ‘Materials and Methods’ section. **c** Graph showing the fractions of RNA bound by IN as a function of IN concentration. Data points (Mean ± SD) derived from four independent experiments. Data were fitted with Kaleida Graph (Synergy software) to Michaelis-Menten equation.

As a control, we studied the interaction of Tat with TAR, the cognate interacting protein. We used a chemically synthesized full-length Tat protein (1-102 aa) to assess the binding with TAR mutants by EMSA assay. Similarly to IN-CTD and consistently with other *in vitro* and *ex vivo* information ^46 47 48 49^, the binding ability of Tat decreased with stem shortening (**Supplementary Fig. 3b**).

Overall, our results suggest that IN-CT interacts with the apical stem-loop of TAR, possibly with its major groove. This is consistent with the CLIP-seq data, which had identified the TAR hexaloop as a major binding site for IN and with a recently published structural model of IN-CTD-ΔCT bound with TAR ^8 50^.

### IN-CTD deeply affect the structure of TAR favoring Tat interaction

In order to extend the binding analysis at the molecular level, the IN-TAR interaction was probed by footprinting techniques. 5’ radiolabeled TAR, alone or complexed to protein, was subjected to partial digestion using RNase T1 that cleaves 3′ to an unpaired guanine. The results shown in **Fig. 3a** indicated that G34 and G36 of TAR were protected in the presence of IN-CTD (**Fig. 3a**, lane 6), instead the binding of IN-CT and IN-CTD-ΔCT did not protect these nucleotides from nuclease digestion (**Fig. 3a**, lanes 4 and 5). This suggested that the CT region is interacting with G34 and G36 nucleotides located in the TAR hexaloop and its junction with the 4-nt stem when CT is part of the whole domain (**Fig. 3a**, lane 6). Surprisingly, we observed prominent cleavage following the residues at positions C41, U38, G32 and, to a lesser extent, G28, induced by the binding of IN-CTD-ΔCT and IN-CTD even in the absence of T1 nuclease (**Supplementary Fig. 4a**, left panel). This reflected the presence of drastic structural constraints on the apical stem-loop of TAR upon the binding of either IN-CTD-ΔCT or IN-CTD, which leads to a spontaneous mechanical breakage. Any nuclease contamination has been observed in protein solution (**Supplementary Fig. 4a**, right panel). Altogether, these digestion patterns showing the protection of G34 and G36 by IN-CTD, but not IN-CTD-ΔCT, as well as the evidence of the structural deformation of TAR induced by the binding of both fragments, indicate that: (i) IN-CTD binds to the TAR apical stem-loop; (ii) IN-CTD modifies the structure of the RNA; (iii) the CT exerts a specific role in the interaction of IN-CTD with the TAR hexaloop and its junction with the stem. The fact that CT binds TAR only in the IN-CTD context, could reflect the necessity for a distortion of TAR by the IN-CTD-ΔCT moiety to allow for the correct binding of CT to the hexaloop. Therefore the role of Tat seems to be that of coating TAR which resulted in a prevention of nuclease digestion (**Fig. 3a**, lane 7).

**Figure 3.**
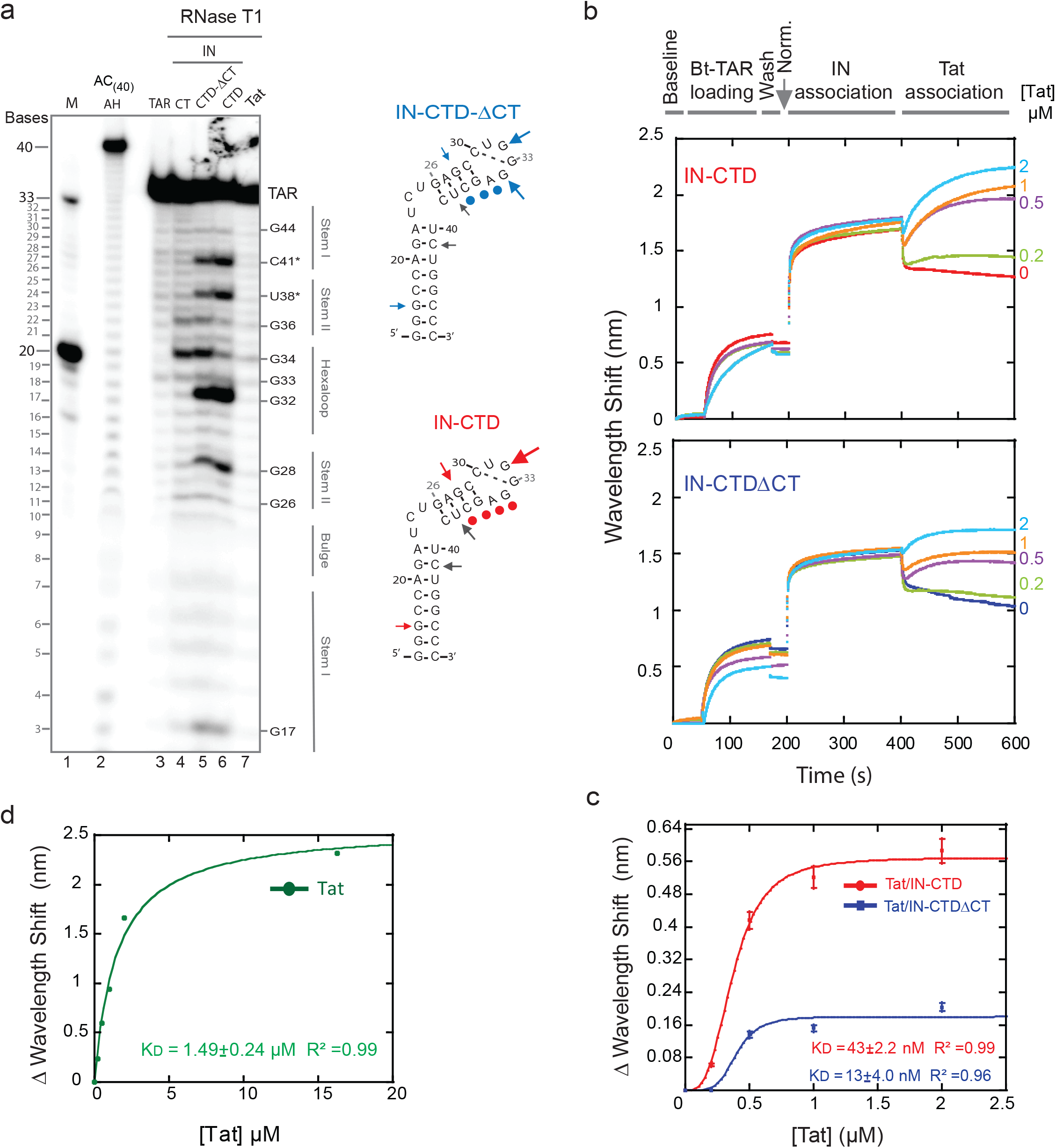
IN induces structural modifications of TAR, promoting Tat binding. **a** Structure probing of the secondary structure of TAR RNA complexed with different fragments of IN and Tat. 5’-end radiolabeled TAR (34-mer) was incubated in the presence or absence of protein for 30 min at 37 °C prior to RNAse T1 treatment, as described in experimental procedures. RNA fragments were separated on a 15% denaturing polyacrylamide sequencing gel. Bands corresponding to certain T1 cleavage (at G bases) products are identified as position markers. Probing gel lanes are as follows: (M) Ladder of two RNA transcripts of 33 and 20 nucleotides in length (lane 1); (AC(40) AH) alkaline ladder of AC_(40)_-mer RNA (lane 2); TAR native (lane3); TAR RNA complexes with IN-CT (lane 4); IN-CTD-ΔCT (lane 5); IN-CTD (lane 6) and Tat (lane 7). Digestion patterns were mapped on TAR secondary structure depicted on the right of the gel: circles identify nucleotides protected from RNAse T1 digestion and arrows mark the RNA cleavage sites. Grey arrows indicated nonspecific hydrolysis. The dimensions of arrows are proportional to the intensity of the band. **b** Real-time measurements of protein-RNA interaction obtained using Interferometry assays (BLItz^®^ System instrument-FortéBio). The 3’ biotinylated TAR (Bt-TAR) was first loaded on the streptavidin-coated biosensor for 100 s (Bt-association) then the unbound RNA was washed for 50 s (baseline). The sensor was absorbed in a solution containing about 90 μM of IN protein for 200 s than incubated with different concentrations of Tat (0.2, 0.5, 1 and 2 μM) for 200 s. **c** Graph showing the wavelength shifts recorded 200 s after Tat addition (at 600 s of kinetics) normalized on the minimum wavelength shifts measured at the moment of Tat addition (400 s), as a function of the corresponding Tat concentrations. Data points were fitted with Hill equation: y= Bmax x^n^ /K_D_+(x ^n^). **d** BLI experiment to measure the affinity of Tat for immobilized Bt-TAR RNA: graph showing the wavelength shifts recorded after 200 s after the start of the protein/RNA binding were plotted against the corresponding Tat concentrations. Data points were fitted with Michaelis-Menten equation and R^2^ and K_D_ values were shown in the graphs inset.

In order to understand whether the structural changes of TAR induced by IN were affecting the binding affinity of Tat, we performed displacement experiments by BLI. 3’ biotinylated TAR was previously immobilized to streptavidin-coated biosensor, then IN association to TAR was monitored as described before. Afterwards, the sensor was absorbed in solutions containing various concentrations of Tat (**Fig. 3b**). We calculated the apparent K_D_ by measuring the Δ Wavelength Shift between the minimum wavelength after Tat injection (around 400 s) and the maximum wavelength reach at 600 s (**Supplementary Fig. 4c**), and we plotted the values against the corresponding Tat concentrations (**Fig. 3c**). The K_D_ of Tat binding to TAR was about 37.5 and 75 times lower in presence of IN-CTD-ΔCT and IN-CTD respectively (**Fig. 3c**) compared to the K_D_ of Tat measured in the same conditions for the naked RNA (**Fig. 3d** and **Supplementary Fig. 4d**). Moreover, the binding kinetics of Tat on IN/TAR complex is consistent with a cooperative interaction (**Fig. 3c**). Notably, the IN-CTD:TAR complex can accommodate more Tat than IN-CTD-ΔCT:TAR, as indicated by the higher B_Max_ (**Fig. 3c**). To exclude that these results derived from a direct interaction between IN-CTD and Tat, we performed a pulldown assay which did not show protein-protein interaction in these conditions (**Supplementary Fig. 4e**).

### Tat competes with IN-CTD and displaces it from TAR

To investigate the mechanism involved in Tat binding to IN:TAR complexes, we performed a dose-dependent competition EMSA (**Fig. 4a**), in the presence of 10 K_D_ IN/TAR ratio. We could confirm that Tat binds to TAR in a dose-response manner (**Fig. 4a**, lanes 13-17). Interestingly, we observed a dose-dependent interference between Tat and IN-CTD for TAR binding at intermediate (**Fig. 4a**, lanes 4 and 5), but not at high concentrations of Tat (**Fig. 4a**, lane 6). Furthermore, no interference was observed at any concentration for IN-CTD-ΔCT (**Fig. 4a**, lanes 9 to 12), confirming that the mechanism was dependent on the presence of the CT tail. Since the migration pattern of the EMSA cannot discriminate between IN-bound and Tat-bound RNAs, we assessed whether the presence of Tat would modify the binding of IN to 3’ biotinylated TAR by pull-down assays (**Fig. 4b**). IN-CTD and IN-CTD-ΔCT were efficiently co-precipitated (**Fig. 4b**, lanes 1 and 5, respectively), whereas the addition of increasing concentrations of Tat serially decreased their binding to TAR (**Fig. 4b**, lanes 2 to 4 and 6 to 8, respectively). Control experiments showed that Tat also interacted with TAR and was not precipitated when biotinylated TAR was absent (**Fig. 4b**, lanes 9 and 10). Altogether these results underline a competition of Tat and IN for TAR binding. The fact that very small differences were detected between IN-CTD and IN-CTD-ΔCT during TAR-pulldown assay (**Fig. 4b**), while they were observed in EMSA (**Fig. 4a**) is likely due to the higher sensitivity offered by the latter, in which nanomolar concentrations of proteins and RNA were used.

**Figure 4.**
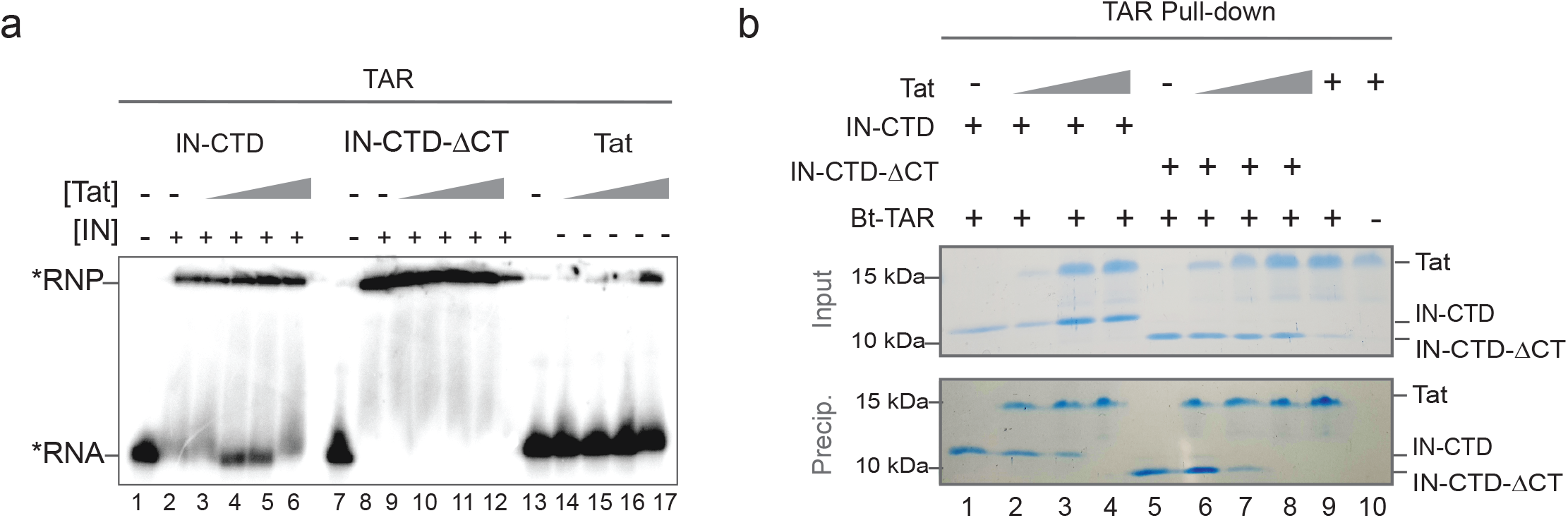
Tat competes with IN-CTD for TAR binding. **a** EMSA assay showing a dose-dependent competition assay of Tat on IN-CTD/TAR or IN-CTD-ΔCT/TAR complexes. Radiolabeled TAR (50 nM) was incubated with 16 fold molar excess of IN-CTD (lanes 2-6) or IN-CTD-ΔCT (lanes 8-12) with or without increasing concentration of Tat (25, 50, 100, 200 nM). Tat *vs* IN-CTD is shown in lanes 3-6 and IN-CTD-ΔCT in lanes 9-12. Control binding of Tat on TAR was also performed using Tat alone (lanes 13-17). **b** Protein co-precipitation with 3′ end-biotinylated TAR (Bt-TAR). IN-CTD (lanes 1-4) or IN-CTD-ΔCT (lanes 5 and 8) were mixed with increasing amount of Tat (1, 2, 3 μg/sample; lanes 2, 3, 4 and 6, 7, 8) and incubated in a buffer containing 200 mM NaCl before co-precipitation. Tat was incubated with Bt-TAR alone (lane 9) or TAR-free beads as a control for unspecific binding (lane 10). Input (20% of total) and pull-down fractions were analyzed by 15% SDS-PAGE followed by Coomassie blue staining.

## DISCUSSION

The C-terminal tail of HIV-1 IN ensures several functions essential for infectivity. A comprehensive understanding of its involvement in the various steps of the infectious cycle is still lacking due to a lack of structural information and to the pleiotropic effects caused by its mutations ^51 21^. Here, we found that this intrinsically disordered region of IN acts as a specific sensor for the peculiar structure of the apical stem-loop of TAR RNA and is directly interacting with its apical hexaloop. We probed IN/TAR interaction by EMSA, nuclease digestion and bio layer interferometry. Data analysis suggested that IN-CTD modified TAR conformation, leading to an enhanced binding of Tat, especially when CT region is present. Moreover, we put forth evidence for an interplay between Tat, IN and TAR, where Tat is competing with IN-CTD for TAR binding and destabilizes preformed IN-CTD:TAR complex.

Mutations of TAR aimed at altering the peculiar structure of the apical portion decreased the relative affinity of IN-CTD compared to IN-CTD-ΔCT, suggesting a role of CT in the recognition of this portion of TAR structure (**Fig. 2**). Importantly, full-length IN presented the same trend of sensitivity for TAR structural mutants (**Supplementary Fig. 3a**). The presence of CT does not seem to be associated with a selective specificity of IN for TAR *in vitro*, as IN-CTD could bind efficiently to other structured RNAs regardless of the presence of CT. In particular, IN-CTD interacts with polyA, DIS, and SD/Ψ RNA elements of HIV-1 5’UTR (**Fig. 1d**). Consistently, the K_D_ measured for IN interaction with all these gRNA elements has been shown to span a narrow range of values ^8^. The better affinity previously observed for the full-length IN ^8^ compared to IN-CTD measured in this work (**Fig. 1f** and **Supplementary Fig. 2b** and **c**) could be due to the presence of additional RNA binding sites within the NTD and CCD domains of full-length IN ^8^ and/or to differences in analytical techniques with respect to those employed here. Thus, the CT region can recognize a TAR RNA with the proper apical stem-loop conformation among possible structural defective TAR conformers, but it is not involved in the discrimination between viral structured RNA regions. Interestingly, this behavior of IN-CT reminds that of p6, the C-terminal domain of HIV-1 Pr55^Gag 52^. Pr55^GagΔp6^ mutant, deleted of the p6 domain, showed no RNA binding specificity compared to the full lengths Pr55^Gag^, suggesting that the presence of this region is required for the specific binding of Pr55^Gag^ to DIS RNA within the 5’UTR of HIV-1 genome ^52^.

Previous biochemical and structural studies have demonstrated that unstructured Tat arginine-rich motif (ARM) penetrates in the TAR major groove made by stem-bulge-stem-loop secondary structures and mostly interacts with the U-rich bulge and nearby double-stranded regions (**Supplementary Fig. 5a**; ^38 53 54 55^). Despite the absence in the CT of a motif comparable to the Arginine stretch of Tat, our results suggest that the unstructured IN-CT region also binds the TAR major groove, but through the interaction with G34 and, to a lesser extent G36, of the hexaloop (**Fig. 3a**). This could explain the lower affinity of IN-CTD for TAR mutants (**Fig. 2**): likely because the altered position of G34 and G36. Interestingly, in a recent report, the IN-CTD:TAR complex has been modeled on the base of the IN-CTD:INI1_183-304_ structure, based on the fact that the same 6 residues were engaged for the binding of IN-CTD to INI1 and TAR (^8,50^ **Supplementary Fig. 5b**). In this model, IN-CTD binds the minor groove of the 4-nt apical stem of TAR. Noteworthy, the last C-terminal modeled residue (D270), immediately preceding the CT tail, is oriented towards the major groove of TAR, which is consistent with our hypothesis (^50^ **Supplementary Fig. 5b**).

The interaction of IN with gRNA has been shown to be critical for the proper localization of vRNP inside the protective capsid lattice ^8,25,26^. In this context, the TAR-selection activity is exerted by IN-CT, while in the Pr160^Gag-Pol^ precursor, could be an additional mechanism to that of the Pr55^Gag^ protein interaction with the packaging signals (reviewed in ^56^) in order to selectively recruit and encaspsidate the HIV-1 gRNA in the viral core.

Our observations show how the interplay between IN, Tat and TAR RNA takes place: first IN-CTD binds TAR through its C-terminal tail by contacting the apical stem-loop and in particular the hexaloop. This interaction modifies the structure of TAR (**Fig. 3a**) favoring Tat binding (**Fig. 3b** and **3c**), which finally displaces IN from the TAR RNA (**Fig. 4**). Noteworthy, the competition of Tat ARM with IN for TAR binding has also been reported elsewhere ^8^. Our data are also fully consistent with recent evidences of the implication of IN in the transcription of the provirus at early times after integration and in a Tat-independent manner ^32^. The authors found that mutation of four lysine residues within the IN-CTD dramatically reduced proviral transcription since all of them are involved in the binding of IN to the viral RNA ^8,26^.

Taken together, these *in vitro* observations suggest a working model in which IN would ensure the very first stages of proviral transcription (**Fig. 5**): IN-CTD, through to its CT tail, selectively binds to the nascent TAR transcript. This binding modifies the structure of TAR, facilitating its interaction with Tat. Therein Tat displaces IN and allows the subsequent transactivation of provirus transcription (**Fig. 5**). This process is dynamic and might also be modulated by post-translational modifications of IN, interaction with cellular partners and/or chromatin remodeling processes.

**Figure 5.**
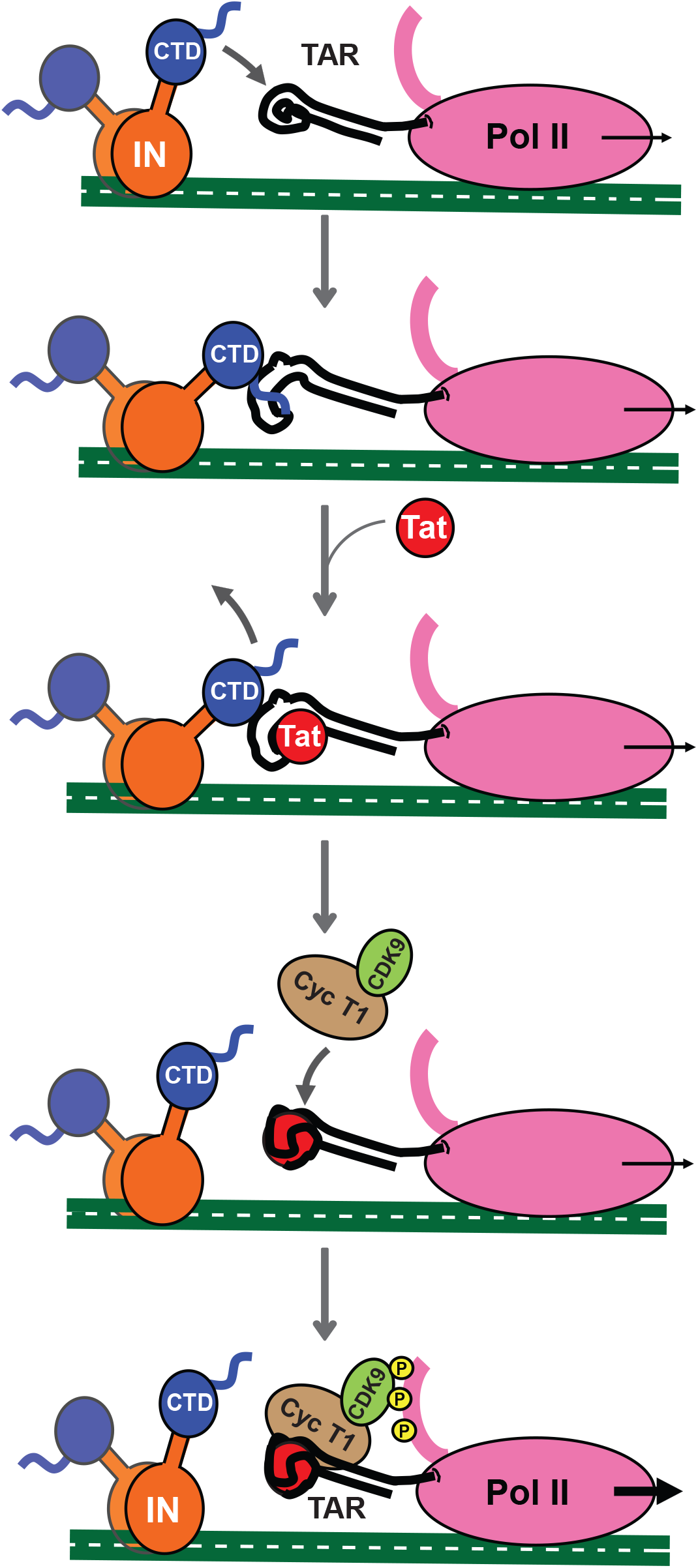
Working Model proposed for the interplay of IN-CTD, Tat and TAR during the early state of HIV-1 proviral transcription. Once the integration has taken place, IN interacts with nascent TAR transcript through its CTD, directly interacting with the major groove and the hexaloop, thanks to its C-terminal tail. The binding induces TAR conformational changes, which promote the interaction of Tat with its substrate. Afterwards, Tat displaces IN-CTD from TAR and recruits the SEC complex in order to boost Pol II transcription.

Interestingly, the structure of the HIV-1 intasome during the strand transfer process, revealed that the inner IN tetrameric core, which contacts both the viral and host DNA, is surrounded by twelve other subunits which display a considerable flexibility ^57^ and might be available for other functions. Unfortunately, the structure of IN after strand transfer is not yet available and the prediction of the presence of CTDs available for transcription is not possible so far.

## METHODS

### Protein expression and purification

Construction of plasmid pET15b, encoding N-terminal 6XHis tagged IN-CTD (aa 220-270) and IN-CTD-ΔCT (aa 220-288) were previously reported ^44^. IN-FL gene from pNL4-3 were cloned in pPROEX-HTa vector in frame with 6XHis tag at N-terminus. All proteins were expressed in *Escherichia coli* BL21(DE3) Rosetta/pLysS strain (Novagen). Cells were grown in LB medium supplemented with 10% (w/v) glucose and protein expression was induced at an optical density at 600 nm (OD_600_) of 0.6 with 1 mM IPTG (isopropyl-β-d-thiogalactopyranoside). Cells were incubated overnight at 18°C under continuous shaking, then pelleted. The cell pellets were resuspended in lysis buffer composed by 50 mM Tris-HCl pH 8, 1 M NaCl, 20 mM imidazole, 0.1 mM EDTA, 2 mM β-mercaptoethanol and 10% (w/v) glycerol. Exclusively for bacterial lysis the buffer was freshly supplemented with 2M urea, 2 mM of Adenosine triphosphate (ATP), 5mM CHAPS (3-[(3-cholamidopropyl)dimethylammonio]-1-propanesulfonate) and 1 tablet of Protease Inhibitor Cocktail (ROCHE, cOmplete™). The preparation was sonicated for 120 s on ice, then the resulting lysate was subjected to centrifugation at 11.000 *g* for 1h. The recovered supernatant was then applied to a HisTrap™ Fast Flow Crude column (Cytiva) and purified by AKTA pure system (Cytiva). The sample was first abundantly washed with lysis buffer containing 100 mM imidazole and 2M NaCl, then the protein was eluted using a gradient up to 500 mM imidazole in 10 column volumes. A second step of purification was carried out using a Superdex 75 10/300 GL column (Cytiva) by an isocratic elution carried out with storing buffer (50 mM HEPES pH 7.5, 1 M NaCl, 5 mM β-mercaptoethanol, and 5% glycerol).

FlagIN-FL (IN-FLm) was expressed in Baby Hamster Kidney suspension cells (BHK21-C13-2P, Sigma-Aldrich) using a vaccinia virus expression system as previously described ^42^.

IN-CT peptide (YGKQMAGDDCVASRQDED) and 101-residue long Tat protein of primary isolate 133 of HIV-1 were chemically synthetized ^58 59^. This Tat has been biochemically characterized and its full biological activity was previously validated ^60^.

### In vitro RNA synthesis, purification, and radiolabeling

We produced several RNAs as listed in Table 1 by using partially double-stranded templates formed by hybridization of T7 promoter-containing DNA oligonucleotides listed in Table 2 and following the protocol detailed in ^61^. Templates for polyA, DIS and SD/Psi RNAs were produced by PCR using pNL4-3 plasmid as template and T7 promoter-containing primers. RNA was transcribed by kit MEGAshortscript™ T7 (Thermo Fisher Scientific) following the manufacturer instructions, de-phosphorylated using Alkaline Phosphatase (New England Biolabs) and purified by phenol:chloroform:isoamyl alcohol (25:24:1) extraction and ethanol precipitation. The 3’ biotinylated TAR and RNA 30-mer (Table 1) were chemically synthesized (Sigma Aldrich). RNA (50 pmol) was radiolabelled at 5′ end with by using 10 units of T4 polynucleotide kinase (New England Biolabs) mixed to 3 μl of γ^32^P-ATP (3000 Ci/mmol 10 mCi/ml, Perkin Elmer) for 1h at 37°C then purified on denaturing 10% (w/v) polyacrylamide gel (29:1) as previously described (Fiorini et al 2012). Before use in binding and structural studies, RNA was heated in refolding buffer (20 mM HEPES pH 7.5, 0.2 M NaCl, 2 mM MgCl_2_, 2 mM DTT) for 3 min at 95°C, followed by 40 min of slow controlled cooling to room temperature, and finally placed on ice.

**Table 1.**
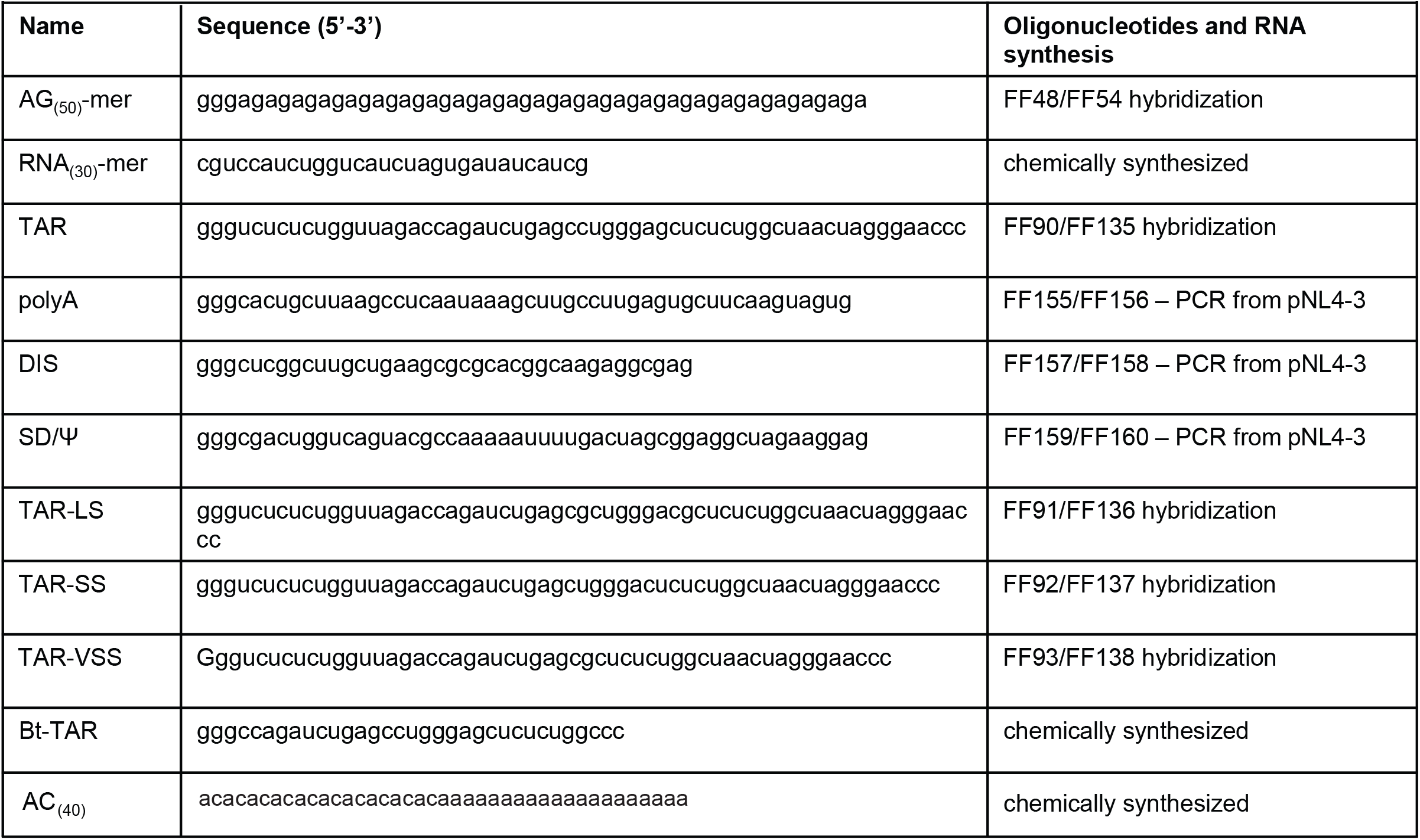
RNAs used in this study.

**Table 2.**
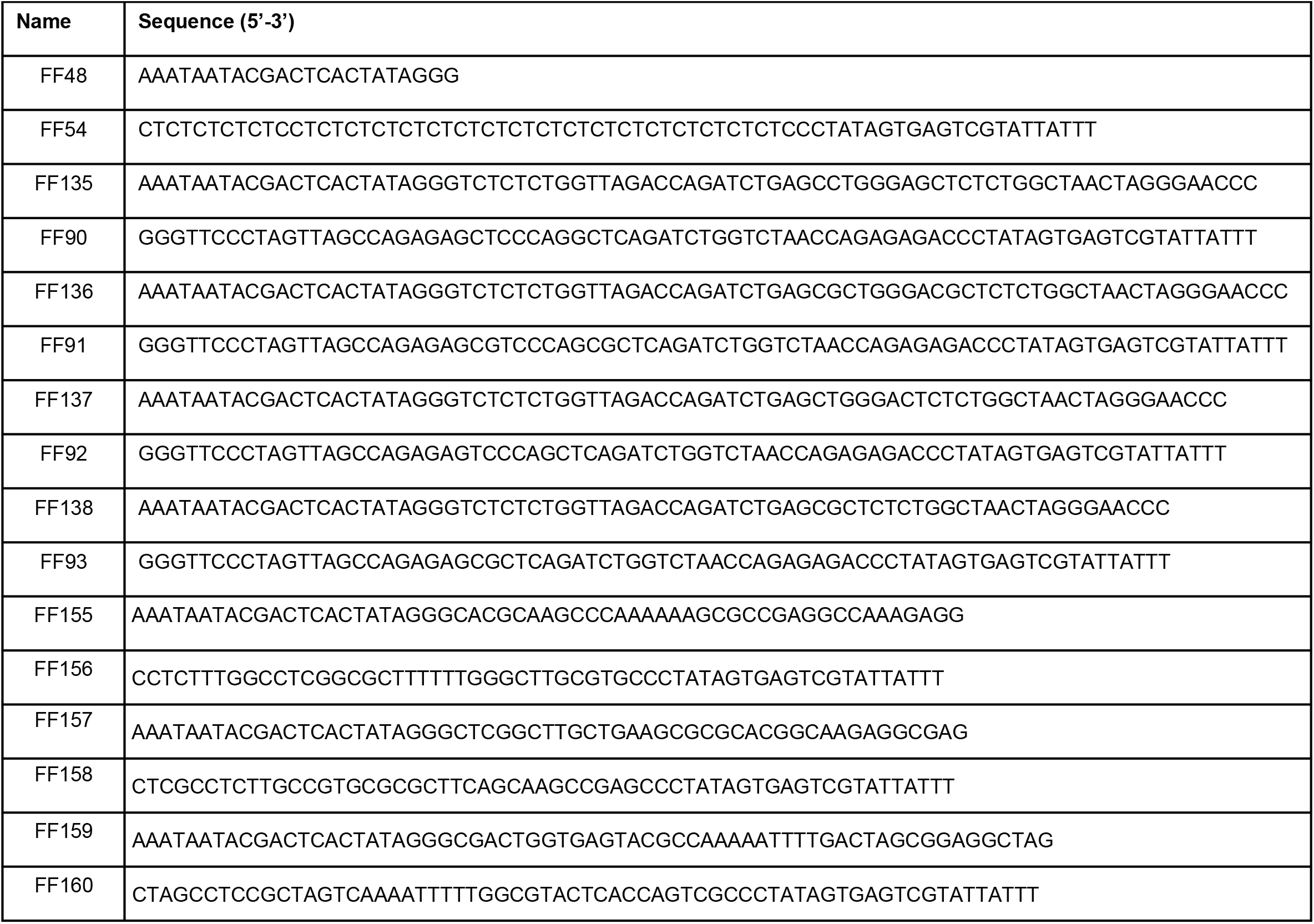
Oligonucleotides used to produce RNA substrates.

### Electrophoretic Mobility Shift Assays (EMSA)

The electrophoretic mobility shift assays (EMSA) were performed as described in ^62^. Samples were prepared by mixing a radiolabelled RNA with increasing concentrations of proteins, as indicated, in a buffer containing 20 mM MES pH 6.0, 150 mM NaCl, 2 mM DTT, 2 mM MgCl_2_, 0.2 μg BSA and 8% (v/v) PEG8000. The samples were incubated at 37 °C for 30 min before being resolved by native 6% polyacrylamide (19:1) gel electrophoresis in 0.5x TAE (Tris acetate EDTA) buffer. Results were analysed by phosphorimaging using ImageQuant software. IN-FLm was incubated for 2h at 4°C with the RNA substrate. For dose-dependent competition assay showed in Fig. 4a, we used in all the samples a constant saturating IN-to-TAR concentrations with an IN-to-TAR ratio per sample exceeding about ten times their respective K_D_. Then, we added increasing concentrations of Tat.

### RNA Structural probing

Enzymatic treatments were performed in 10 μl of reaction mix containing 0.5 pmol of 5’ radiolabeled RNA, 0.2 μg of yeast tRNA, 1 × Structure buffer (Thermo Fisher Scientific) and 0.01 U of RNase T1 (Thermo Fisher Scientific). Incubation was done at 37°C for 5 min. Reactions were stopped by addition of 40 μl of quenching buffer composed by 10 mM HEPES pH 7.5, 1 mM EDTA and 3% SDS. Partial alkaline hydrolysis was performed as follows: 10 μl of reaction mix containing 0.2 pmol of RNA, 1 μg of yeast tRNA, 1× Alkaline Hydrolysis buffer, were incubated at 95°C for 12 min then quenched wih 2x denaturing loading buffer and placed on ice and. For RNA/protein complexes, 0.5 pmol RNA was previously incubated with 36 pmol of protein at 37°C for 30 min then treated with RNase T1. After quenching, samples were phenol extracted and ethanol precipitated. After recovery from precipitation, all samples were run on a 15% sequencing polyacrylamide gel in 0.5 × TBE. (Tris Borate EDTA). Results were analysed by phosphorimaging.

### Pulldown assay

The pulldown assays were conducted as described before ^61^. Briefly, proteins were mixed in binding buffer (40 mM HEPES pH 7.5, 2.5 mM MgCl_2_, 2mM β-mercaptoethanol and 5% glycerol) adjusting the final NaCl concentration to 150 mM. Samples were complemented or not with 600 pmol of 3′ end–biotinylated TAR (Table 1) in a final volume of 30 μl, and incubated 30 min at 37 °C. To the mix were added 8 μl of magnetic streptavidin beads (Dynabeads MyOne, Thermo Fischer Scientific) and further incubated for 1h at 4°C in gentle rotation. The resin was washed three times with 500 μl BB-200 (40 mM HEPES pH 7.5, 200 mM NaCl, 2.5 mM MgCl_2_, 2mM β-mercaptoethanol and 10% glycerol) on ice and proteins were eluted with SDS loading buffer and analysed on polyacrylamide 16% (37.5:1) SDS-PAGE. For Histidine pulldown assay, magnetic streptavidin resin was replaced with HisPur™ Ni-NTA Magnetic Beads and protein elution was done with 0.5 M –Imidazole containing buffer.

### Bio-Layer interferometry

For Bio-Layer Interferometry (BLI) analysis we used the BLItz platform (FortéBio). For high throughput experiments such as shown in Fig. 1 and Supplementary Fig. 2e we used the Octet RED96e System (FortéBio). All Measurements were performed in assay buffer composed of 50 mM Hepes pH 7.5, 200 mM NaCl, 2 mM β-mercaptoethanol, 5% glycerol, 2 mM MgCl_2_, 1 μM ZnSO_4_, 0.5 M BSA. For BLItz platform we used Streptavidin (SA) Biosensors (ForteBio) and for Octet RED96e the Streptavidin (SAX) Biosensors (FortéBio) that were hydrated for 10 min in assay buffer then plunged in a solution containing 1 μM 3′ end–biotinylated TAR RNA in assay buffer with 1x RNAase inhibitor (RNA Secure, Invitrogen) and 0.1 μM BSA for the RNA loading step. A wash was performed after RNA loading. In BLItz experiment the association of the protein to RNA was monitored in real-time for 200 s: the hydrated biosensor tip was placed in a 1.5 ml black assay tube containing 200 μl of protein solution in assay buffer as indicated. Afterward, a dissociation step or a second association with Tat, was performed for 200 s. Biosensors were discarded after each measurement. All kinetic assays showed in Fig. 1 and Supplementary Fig. 2e were performed using Octet RED96e system and carried out using black 96-well plates and samples were diluted in freshly prepared assay buffer and incubated at 37°C with an orbital shake. Association and dissociation steps were as in the BLItz experiments. Each time reference sensors and negative control sensors were included. Sensorgrams were exported and data analysis was performed with Kaleida Graph software (Synergy Software).

## Acknowledgements

This work was supported by the French Agency for Research on AIDS and Viral Hepatitis (ECTZ72240-1 to F.F., M.R. and M.N.; PhD fellowship for C.R.). We acknowledge: the support and the use of resources of the French Infrastructure for Integrated Structural Biology FRISBI ANR-10-INBS-05 and of Instruct-ERIC; Sylvie Ricard-Blum and the Interaction platform of the UMR 5246 ICBMS for the experiments on the Octet RED96e; the Protein Science Facility (PSF) of SFR Biosciences (UAR3444/CNRS, US8/Inserm, ENS de Lyon, UCBL) especially Virginie Gueguen-Chaignon and Eric Desis for their assistance, and Frédéric Galisson for the help with radioactive sources. We are grateful to the Viral DNA Integration and Chromatin Dynamics Network (DyNAVir) for fruitful discussions and to Anne-Catherine Dock-Bregeon for critical reading of the manuscript. We acknowledge Andrea Cimarelli for the pNL4-3 plasmid donation.

## Author Contributions

C.R., C.L., and F.F. conceived and designed experiments. C.R., C.L., A.E.M., C.G., L.B. D.L. and F.F. performed the experiments and/or analyzed data. J.B. and M.R. prepared protein from mammalian expression system; C.R., C.L., A.E.M., C.G., L.B. D.L, M.N., M.R., P.G and. F.F. discussed the results; F.F. wrote the paper. All authors read and edited the manuscript.

## References

1. Castello, A. et al. Insights into RNA biology from an atlas of mammalian mRNA-binding proteins. Cell 149, 1393–406 (2012).

2. Beckmann, B.M., Castello, A. & Medenbach, J. The expanding universe of ribonucleoproteins: of novel RNA-binding proteins and unconventional interactions. Pflugers Arch 468, 1029–40 (2016).

3. Hentze, M.W., Castello, A., Schwarzl, T. & Preiss, T. A brave new world of RNA-binding proteins. Nat Rev Mol Cell Biol 19, 327–341 (2018).

4. Cassiday, L.A. & Maher, L.J., 3rd. Having it both ways: transcription factors that bind DNA and RNA. Nucleic Acids Res 30, 4118–26 (2002).

5. Brodsky, S. et al. Intrinsically Disordered Regions Direct Transcription Factor In Vivo Binding Specificity. Mol Cell 79, 459–471 e4 (2020).

6. Garcia-Moreno, M., Jarvelin, A.I. & Castello, A. Unconventional RNA-binding proteins step into the virus-host battlefront. Wiley Interdiscip Rev RNA 9, e1498 (2018).

7. Basu, S. & Bahadur, R.P. A structural perspective of RNA recognition by intrinsically disordered proteins. Cell Mol Life Sci 73, 4075–84 (2016).

8. Kessl, J.J. et al. HIV-1 Integrase Binds the Viral RNA Genome and Is Essential during Virion Morphogenesis. Cell 166, 1257–1268 e12 (2016).

9. Maertens, G.N., Engelman, A.N. & Cherepanov, P. Structure and function of retroviral integrase. Nat Rev Microbiol (2021).

10. Sakai, H. et al. Integration is essential for efficient gene expression of human immunodeficiency virus type 1. J Virol 67, 1169–74 (1993).

11. Esposito, D. & Craigie, R. HIV integrase structure and function. Adv Virus Res 52, 319–33 (1999).

12. Dyda, F. et al. Crystal structure of the catalytic domain of HIV-1 integrase: similarity to other polynucleotidyl transferases. Science 266, 1981–6 (1994).

13. Zheng, R., Jenkins, T.M. & Craigie, R. Zinc folds the N-terminal domain of HIV-1 integrase, promotes multimerization, and enhances catalytic activity. Proc Natl Acad Sci U S A 93, 13659–64 (1996).

14. Cai, M. et al. Solution structure of the N-terminal zinc binding domain of HIV-1 integrase. Nat Struct Biol 4, 567–77 (1997).

15. Lee, S.P., Xiao, J., Knutson, J.R., Lewis, M.S. & Han, M.K. Zn2+ promotes the self-association of human immunodeficiency virus type-1 integrase in vitro. Biochemistry 36, 173–80 (1997).

16. Eijkelenboom, A.P. et al. The DNA-binding domain of HIV-1 integrase has an SH3-like fold. Nat Struct Biol 2, 807–10 (1995).

17. Woerner, A.M. & Marcus-Sekura, C.J. Characterization of a DNA binding domain in the C-terminus of HIV-1 integrase by deletion mutagenesis. Nucleic Acids Res 21, 3507–11 (1993).

18. Engelman, A., Hickman, A.B. & Craigie, R. The core and carboxyl-terminal domains of the integrase protein of human immunodeficiency virus type 1 each contribute to nonspecific DNA binding. J Virol 68, 5911–7 (1994).

19. Jenkins, T.M., Engelman, A., Ghirlando, R. & Craigie, R. A soluble active mutant of HIV-1 integrase: involvement of both the core and carboxyl-terminal domains in multimerization. J Biol Chem 271, 7712–8 (1996).

20. Lutzke, R.A., Vink, C. & Plasterk, R.H. Characterization of the minimal DNA-binding domain of the HIV integrase protein. Nucleic Acids Res 22, 4125–31 (1994).

21. Dar, M.J. et al. Biochemical and virological analysis of the 18-residue C-terminal tail of HIV-1 integrase. Retrovirology 6, 94 (2009).

22. De Houwer, S. et al. Identification of residues in the C-terminal domain of HIV-1 integrase that mediate binding to the transportin-SR2 protein. J Biol Chem 287, 34059–68 (2012).

23. Mauro, E. et al. Human H4 tail stimulates HIV-1 integration through binding to the carboxy-terminal domain of integrase. Nucleic Acids Res 47, 3607–3618 (2019).

24. Vink, C., Oude Groeneger, A.M. & Plasterk, R.H. Identification of the catalytic and DNA-binding region of the human immunodeficiency virus type I integrase protein. Nucleic Acids Res 21, 1419–25 (1993).

25. Madison, M.K. et al. Allosteric HIV-1 Integrase Inhibitors Lead to Premature Degradation of the Viral RNA Genome and Integrase in Target Cells. J Virol 91(2017).

26. Elliott, J.L. et al. Integrase-RNA interactions underscore the critical role of integrase in HIV-1 virion morphogenesis. Elife 9(2020).

27. Elliott, J.L. & Kutluay, S.B. Going beyond Integration: The Emerging Role of HIV-1 Integrase in Virion Morphogenesis. Viruses 12(2020).

28. Fontana, J. et al. Distribution and Redistribution of HIV-1 Nucleocapsid Protein in Immature, Mature, and Integrase-Inhibited Virions: a Role for Integrase in Maturation. J Virol 89, 9765–80 (2015).

29. Cereseto, A. et al. Acetylation of HIV-1 integrase by p300 regulates viral integration. EMBO J 24, 3070–81 (2005).

30. Manganaro, L. et al. Concerted action of cellular JNK and Pin1 restricts HIV-1 genome integration to activated CD4+ T lymphocytes. Nat Med 16, 329–33 (2010).

31. Zamborlini, A. et al. Impairment of human immunodeficiency virus type-1 integrase SUMOylation correlates with an early replication defect. J Biol Chem 286, 21013–22 (2011).

32. Winans, S. & Goff, S.P. Mutations altering acetylated residues in the CTD of HIV-1 integrase cause defects in proviral transcription at early times after integration of viral DNA. PLoS Pathog 16, e1009147 (2020).

33. Jonkers, I. & Lis, J.T. Getting up to speed with transcription elongation by RNA polymerase II. Nat Rev Mol Cell Biol 16, 167–77 (2015).

34. Mousseau, G. & Valente, S.T. Role of Host Factors on the Regulation of Tat-Mediated HIV-1 Transcription. Curr Pharm Des 23, 4079–4090 (2017).

35. Schulze-Gahmen, U. & Hurley, J.H. Structural mechanism for HIV-1 TAR loop recognition by Tat and the super elongation complex. Proc Natl Acad Sci U S A 115, 12973–12978 (2018).

36. Sobhian, B. et al. HIV-1 Tat assembles a multifunctional transcription elongation complex and stably associates with the 7SK snRNP. Mol Cell 38, 439–51 (2010).

37. He, N. et al. HIV-1 Tat and host AFF4 recruit two transcription elongation factors into a bifunctional complex for coordinated activation of HIV-1 transcription. Mol Cell 38, 428–38 (2010).

38. Pham, V.V. et al. HIV-1 Tat interactions with cellular 7SK and viral TAR RNAs identifies dual structural mimicry. Nat Commun 9, 4266 (2018).

39. Wei, P., Garber, M.E., Fang, S.M., Fischer, W.H. & Jones, K.A. A novel CDK9-associated C-type cyclin interacts directly with HIV-1 Tat and mediates its high-affinity, loop-specific binding to TAR RNA. Cell 92, 451–62 (1998).

40. Ne, E., Palstra, R.J. & Mahmoudi, T. Transcription: Insights From the HIV-1 Promoter. Int Rev Cell Mol Biol 335, 191–243 (2018).

41. Craigie, R. The molecular biology of HIV integrase. Future Virol 7, 679–686 (2012).

42. Levy, N. et al. Production of unstable proteins through the formation of stable core complexes. Nat Commun 7, 10932 (2016).

43. Eijkelenboom, A.P. et al. Refined solution structure of the C-terminal DNA-binding domain of human immunovirus-1 integrase. Proteins 36, 556–64 (1999).

44. Kanja, M. et al. NKNK: a New Essential Motif in the C-Terminal Domain of HIV-1 Group M Integrases. J Virol 94(2020).

45. Lodi, P.J. et al. Solution structure of the DNA binding domain of HIV-1 integrase. Biochemistry 34, 9826–33 (1995).

46. Berkhout, B. & Jeang, K.T. Detailed mutational analysis of TAR RNA: critical spacing between the bulge and loop recognition domains. Nucleic Acids Res 19, 6169–76 (1991).

47. Churcher, M.J. et al. High affinity binding of TAR RNA by the human immunodeficiency virus type-1 tat protein requires base-pairs in the RNA stem and amino acid residues flanking the basic region. J Mol Biol 230, 90–110 (1993).

48. Roy, S., Delling, U., Chen, C.H., Rosen, C.A. & Sonenberg, N. A bulge structure in HIV-1 TAR RNA is required for Tat binding and Tat-mediated trans-activation. Genes Dev 4, 1365–73 (1990).

49. Delling, U. et al. Conserved nucleotides in the TAR RNA stem of human immunodeficiency virus type 1 are critical for Tat binding and trans activation: model for TAR RNA tertiary structure. J Virol 66, 3018–25 (1992).

50. Dixit, U. et al. INI1/SMARCB1 Rpt1 domain mimics TAR RNA in binding to integrase to facilitate HIV-1 replication. Nat Commun 12, 2743 (2021).

51. Lu, R., Ghory, H.Z. & Engelman, A. Genetic analyses of conserved residues in the carboxyl-terminal domain of human immunodeficiency virus type 1 integrase. J Virol 79, 10356–68 (2005).

52. Dubois, N. et al. The C-terminal p6 domain of the HIV-1 Pr55(Gag) precursor is required for specific binding to the genomic RNA. RNA Biol 15, 923–936 (2018).

53. Puglisi, J.D., Tan, R., Calnan, B.J., Frankel, A.D. & Williamson, J.R. Conformation of the TAR RNA-arginine complex by NMR spectroscopy. Science 257, 76–80 (1992).

54. Weeks, K.M., Ampe, C., Schultz, S.C., Steitz, T.A. & Crothers, D.M. Fragments of the HIV-1 Tat protein specifically bind TAR RNA. Science 249, 1281–5 (1990).

55. Cordingley, M.G. et al. Sequence-specific interaction of Tat protein and Tat peptides with the transactivation-responsive sequence element of human immunodeficiency virus type 1 in vitro. Proc Natl Acad Sci U S A 87, 8985–9 (1990).

56. Comas-Garcia, M., Davis, S.R. & Rein, A. On the Selective Packaging of Genomic RNA by HIV-1. Viruses 8(2016).

57. Passos, D.O. et al. Cryo-EM structures and atomic model of the HIV-1 strand transfer complex intasome. Science 355, 89–92 (2017).

58. Guillon, C., Stankovic, K., Ataman-Onal, Y., Biron, F. & Verrier, B. Evidence for CTL-mediated selection of Tat and Rev mutants after the onset of the asymptomatic period during HIV type 1 infection. AIDS Res Hum Retroviruses 22, 1283–92 (2006).

59. Mayol, K., Munier, S., Beck, A., Verrier, B. & Guillon, C. Design and characterization of an HIV-1 Tat mutant: inactivation of viral and cellular functions but not antigenicity. Vaccine 25, 6047–60 (2007).

60. Foucault, M. et al. UV and X-ray structural studies of a 101-residue long Tat protein from a HIV-1 primary isolate and of its mutated, detoxified, vaccine candidate. Proteins 78, 1441–56 (2010).

61. Fiorini, F., Bonneau, F. & Le Hir, H. Biochemical characterization of the RNA helicase UPF1 involved in nonsense-mediated mRNA decay. Methods Enzymol 511, 255–74 (2012).

62. Fiorini, F., Boudvillain, M. & Le Hir, H. Tight intramolecular regulation of the human Upf1 helicase by its N-and C-terminal domains. Nucleic Acids Res 41, 2404–15 (2013).

63. Gruber, A.R., Lorenz, R., Bernhart, S.H., Neubock, R. & Hofacker, I.L. The Vienna RNA websuite. Nucleic Acids Res 36, W70–4 (2008).

64. Lorenz, R. et al. ViennaRNA Package 2.0. Algorithms Mol Biol 6, 26 (2011).

65. Robert, X. & Gouet, P. Deciphering key features in protein structures with the new ENDscript server. Nucleic Acids Res 42, W320–4 (2014).

66. Muller-Esparza, H., Osorio-Valeriano, M., Steube, N., Thanbichler, M. & Randau, L. Bio-Layer Interferometry Analysis of the Target Binding Activity of CRISPR-Cas Effector Complexes. Front Mol Biosci 7, 98 (2020).

