## Supplementary Figures for "The HIV-1 Integrase C-Terminal domain induces TAR RNA structural changes promoting Tat binding"

### Supplementary Figure 1

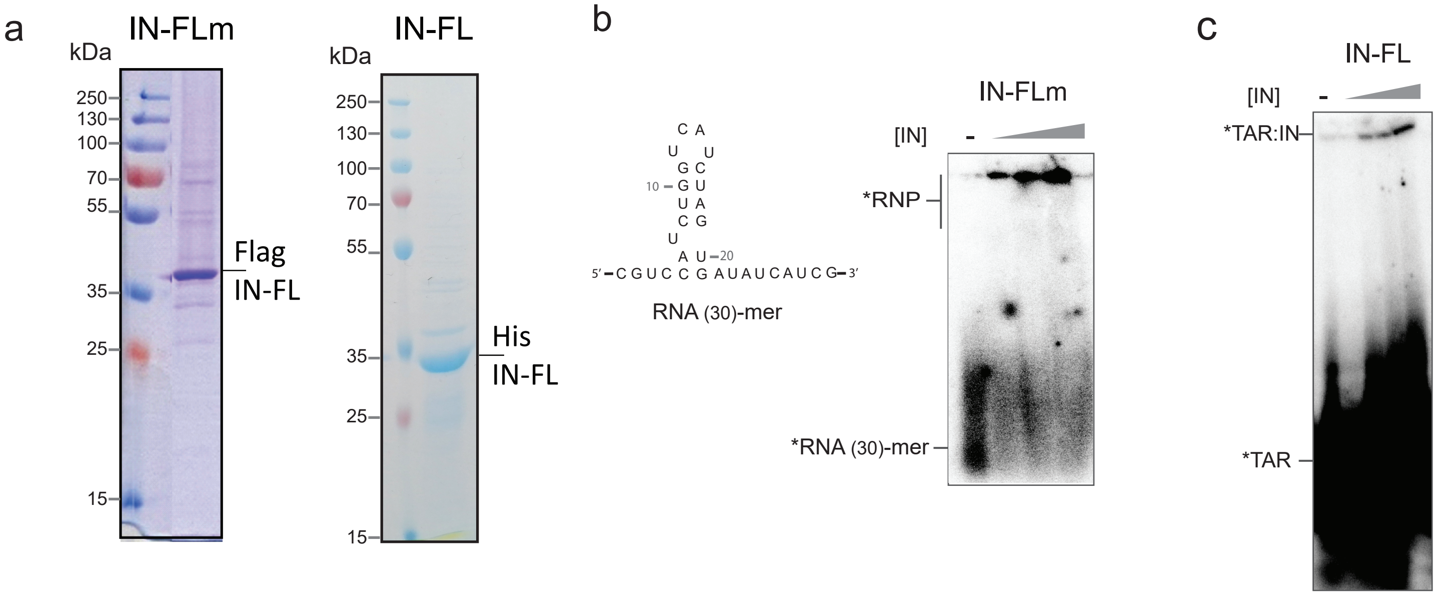

### Supplementary Figure 2

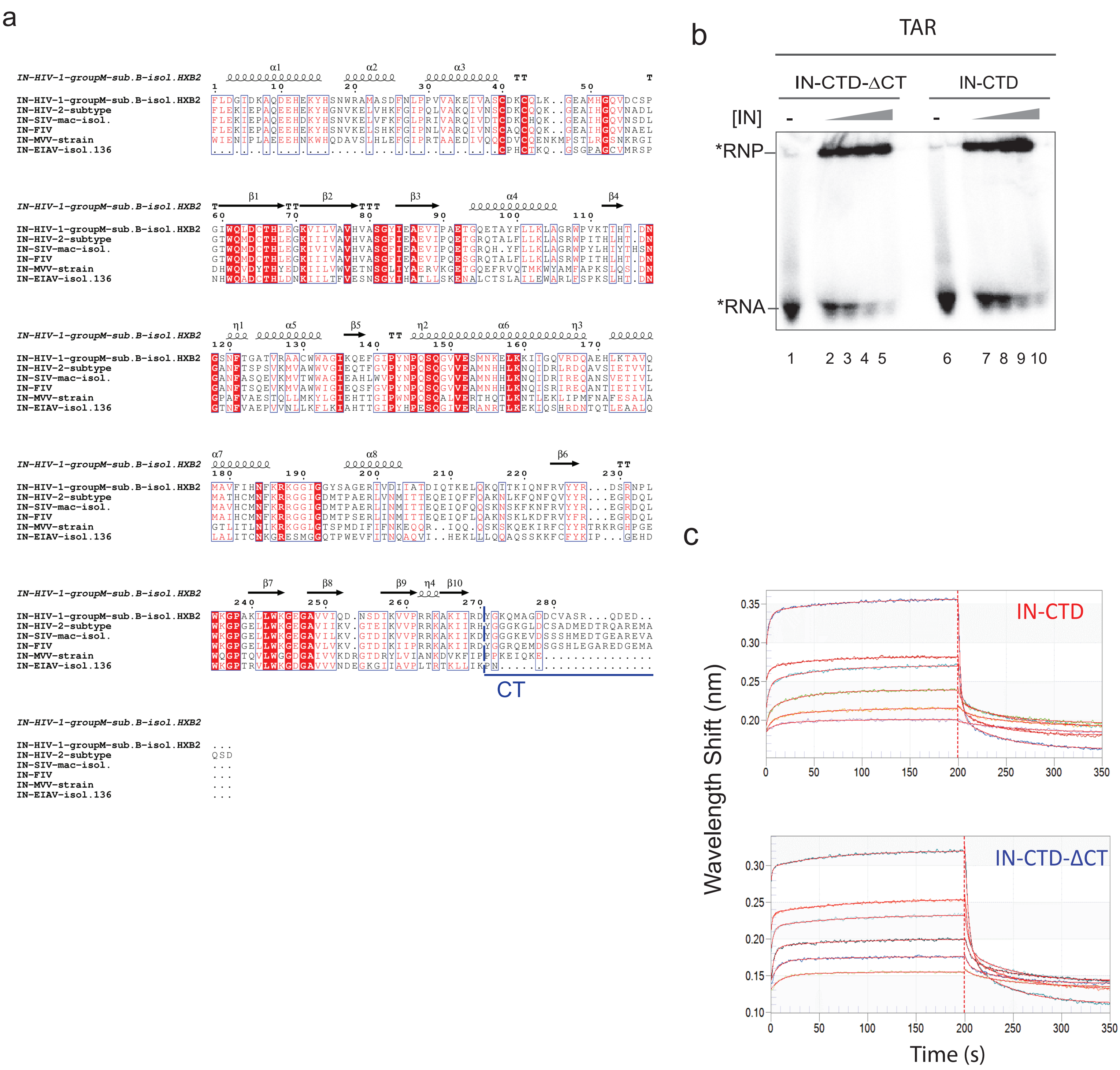

### Supplementary Figure 3

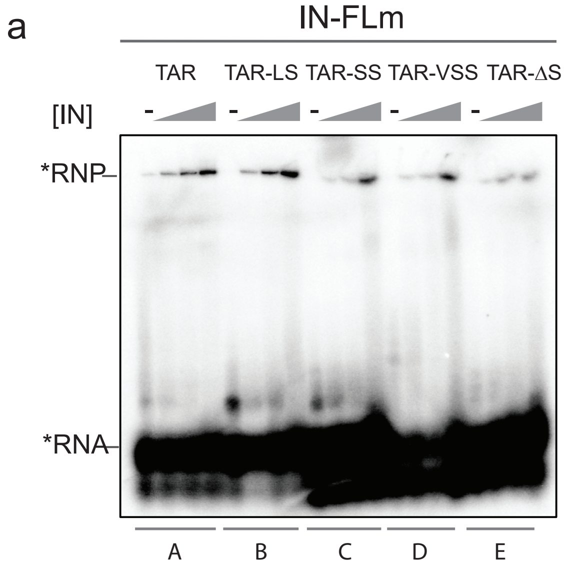


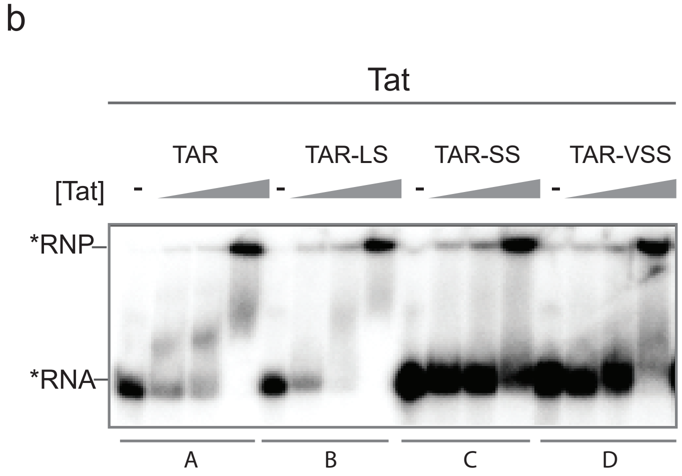

### Supplementary Figure 4

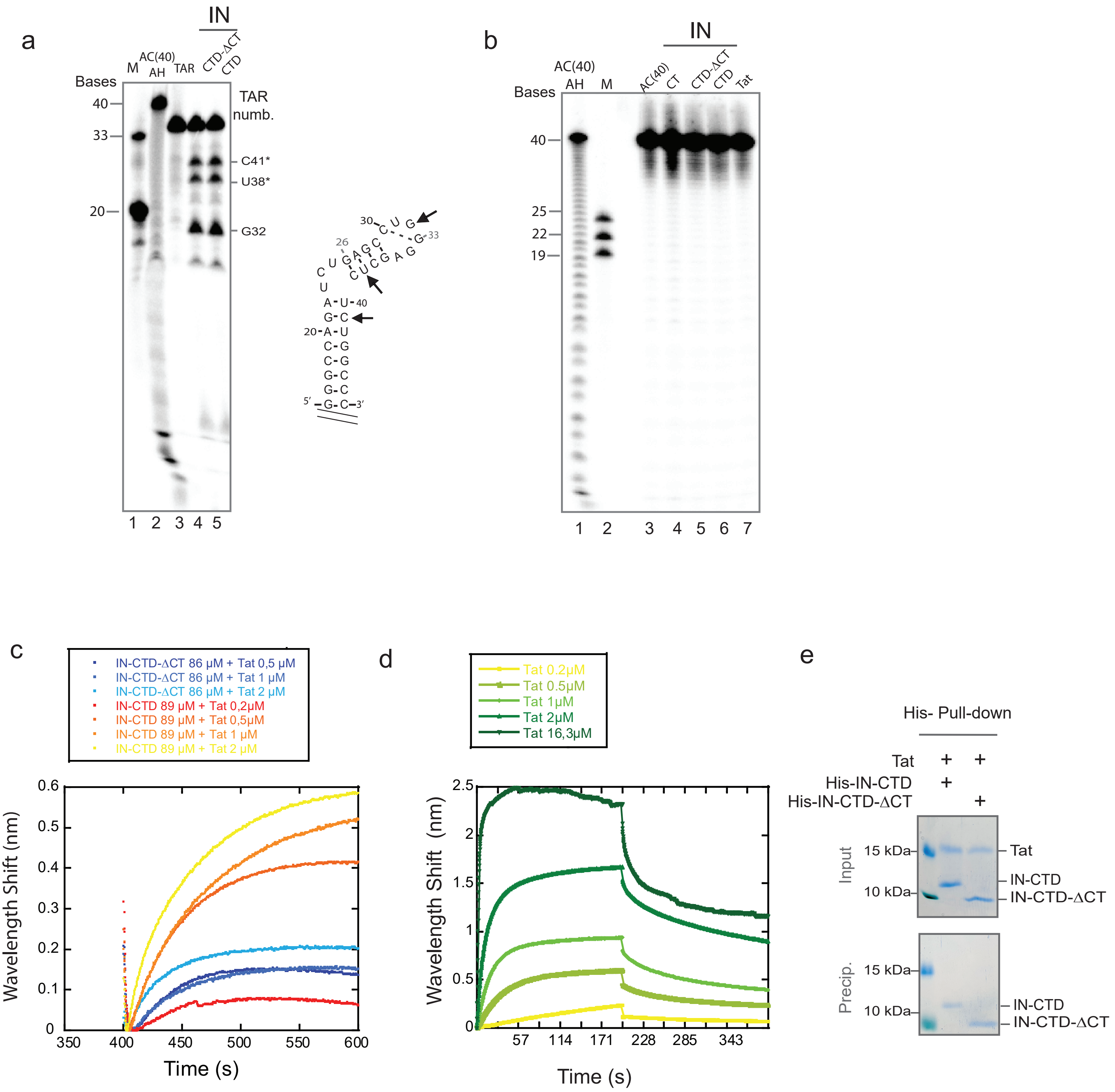

### Supplementary Figure 5

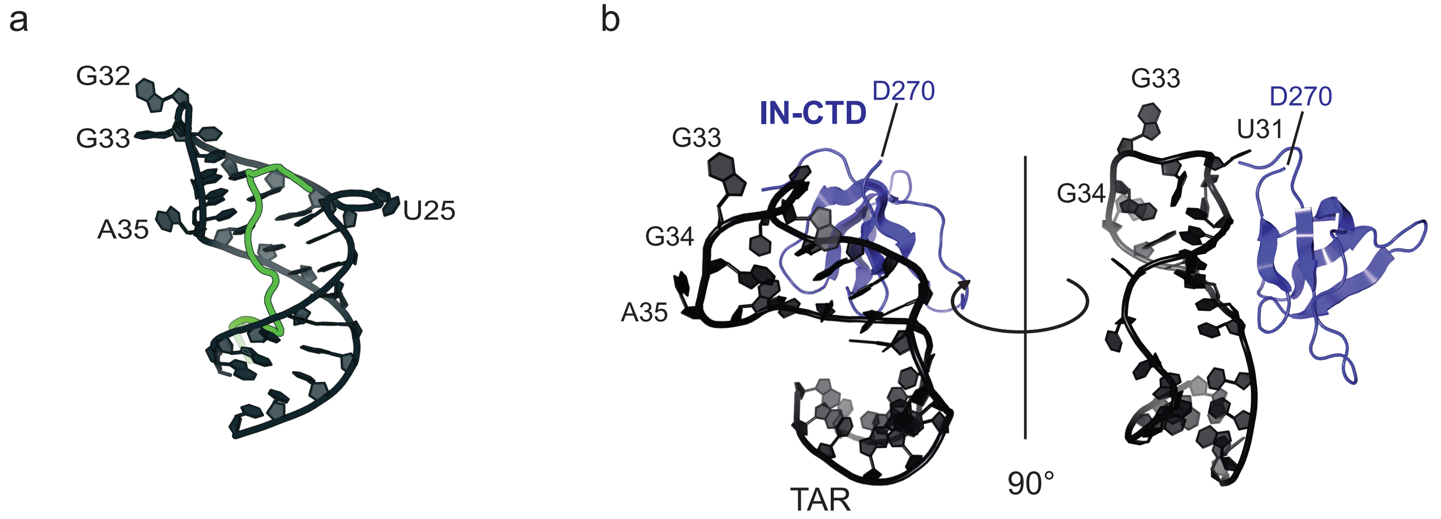


Tat(44-60)-TAR-HIV1
PDB 6MCE
(Pham et al. 2018)
GISYGRKKRRQRRRAHQ

model from Dixit U, Bhutoria S et al. 2021

**Supplementary Figure 1. a** SDS-PAGE illustrating flag-tagged IN-FL expressed from mammalian expression system and IN-FL from *E. coli* used in this study. **b** Structural model (left panel) of a weakly structured RNA 30-mer used for EMSA assay (right panel) illustrating the interaction of IN-FLm. The RNA substrate was incubated with increasing concentrations of proteins (0; 100; 200; 400 μM) for 30 minutes at 37°C in binding buffer as indicated in experimental procedures. **c** EMSA assay indicating the binding of IN-FL produced in *E.coli* to TAR substrate.
